# The Architecture of Sponge Choanocyte Chambers Maximizes Mechanical Pumping Efficiency

**DOI:** 10.1101/2024.02.22.581376

**Authors:** Takumi Ogawa, Shuji Koyama, Toshihiro Omori, Kenji Kikuchi, Hélène de Maleprade, Raymond E. Goldstein, Takuji Ishikawa

## Abstract

Sponges, the basalmost members of the animal kingdom, exhibit a range of complex architectures in which microfluidic channels connect multitudes of spherical chambers lined with choanocytes, flagellated filter-feeding cells. Choanocyte chambers can possess scores or even hundreds of such cells, which drive complex flows entering through porous walls and exiting into the sponge channels. One of the mysteries of the choanocyte chamber is its spherical shape, as it seems inappropriate for inducing directional transport since many choanocyte flagella beat in opposition to such a flow. Here we combine direct imaging of choanocyte chambers in living sponges with computational studies of many-flagella models to understand the connection between chamber architecture and directional flow. We find that those flagella that beat against the flow play a key role in raising the pressure inside the choanocyte chamber, with the result that the mechanical pumping efficiency, calculated from the pressure rise and flow rate, reaches a maximum at a small outlet opening angle. Comparison between experimental observations and the results of numerical simulations reveal that the chamber diameter, flagellar wave number and the outlet opening angle of the freshwater sponge *E. muelleri*, as well as several other species, are related in a manner that maximizes the mechanical pumping efficiency. These results indicate the subtle balances at play during morphogenesis of choanocyte chambers, and give insights into the physiology and body design of sponges.

## I. INTRODUCTION

Sponges are among the oldest multicellular animals, with sponge-like fossils having been found that date as far back as 550-760 million years [1]. As filter feeders with a distinct pump-filter apparatus, sponges (phylum Porifera) can process several hundred times their body volume of water per hour [2, 3]. Because of this, they play a significant role in nutrient cycling within marine ecosystems such as coral reefs [4–6]. To achieve this high-performance filtering, sponges have an evolved aquiferous system that allows water to flow through their bodies [7, 8], and are divided into several classes (termed *asconoid, syconoid*, and *leuconoid*) according to the level of complexity of this internal microfluidic system [9]. Leuconoid sponges have the most complex architecture, composed of incurrent and excurrent canals. Water entering their body through the inlets (ostia) on the surface of the sponge body is carried through the incurrent/excurrent canals and exits from the outlet (osculum), as shown in Fig. 1 for the case of the freshwater sponge *Ephydatia muelleri*. The spherical chambers act as pumps between the incurrent and excurrent canals, and are lined with flagellated cells termed choanocytes arranged in a radial pattern with flagella directed toward the center of the sphere, supporting traveling waves of motion that direct flow toward the center. Filtering of incoming water is achieved by a collar of microvilli anchored near the apex from which emanates the flagellum of each cell.

**FIG. 1.**
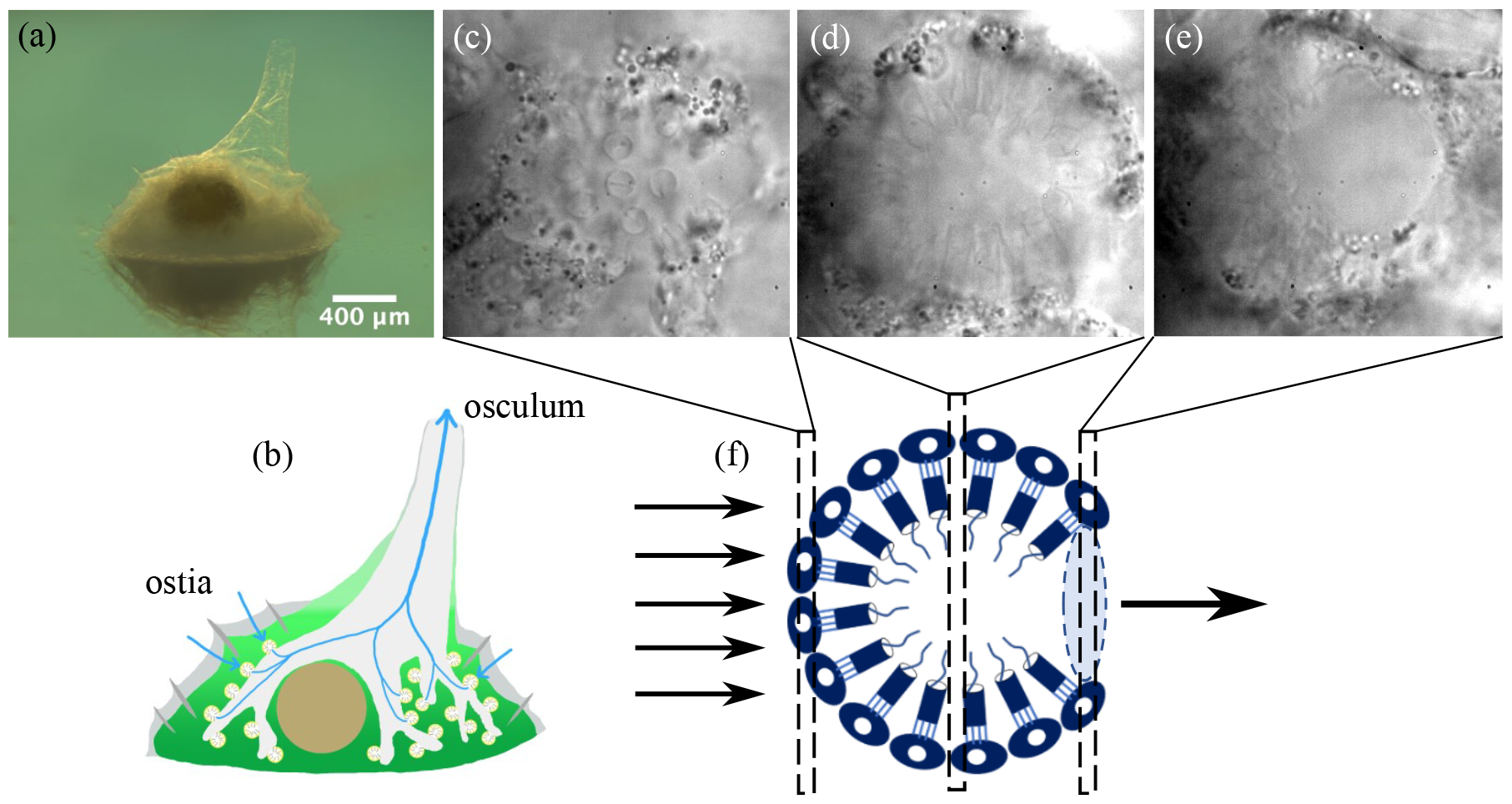
The fresh water sponge *E. muelleri*. (a) Stereomicroscopic image, with the body shown reflected in glass. (b) Schematic diagram of the aquiferous canal system. Blue lines indicate the direction of water flow. (c-e) Microscopic images of the choanocyte chamber at the bottom, middle and top focal planes as indicated in the schematic (f). Parallel arrows indicate fluid flow into a choanocyte chamber, while blue shaded ellipse indicates location of the opening from which fluid exits.

Both phylogenomics [10–12] and morphology [13, 14] support the “Porifera-first” theory of animal evolution. While some studies suggest that the common ancestor of Metazoa appeared from a unicellular choanoflagellate ancestor, because choanoflagellates are closely related to Metazoa and are morphologically similar to sponge choanocytes [13, 15, 16]—it has been confirmed from a molecular phylogenetic view that the two groups are monophyletic [17] and it is difficult to assume homology between these two cells due to functional differences [18]. Thus, while the analysis of the evolution from unicellular to multicellular life remains unsettled, it remains likely that sponges are the oldest animals, and their simple body plans make them a model organism for studies of both morphogenesis and physiology.

The spherical arrangement of choanocytes (Fig. 1(c-f) appears at first glance to be ill-suited to yield directional flow through the chamber, since inevitably the flagella of some of the choanocytes would beat in opposition to the flow. Since sponges have survived from the distant past, it is natural to hypothesize that this spherical shape confers an evolutionary benefit. Here we seek to understand this issue from the perspective of fluid mechanics.

Vogel’s pioneering early work was the first to highlight the fluid mechanics of sponges [19–21]. While the properties of the “sponge pump” have been of interest since then [22, 23], they remain incompletely understood [24–26]. One of the most important issues is the connection between large-scale flows external to the sponge and the flows within. Although the sizes of sponges span an enormous range, even taking a modest vertical scale *L* ∼ 10 − 10^2^ cm and the typical ambient flow speeds *U* ∼ 10 − 10^2^ cm/s the Reynolds number *Re* = *UL/ν* in water is 10^4^ − 10^5^ outside a sponge. With such a large *Re* we expect a viscous boundary to form at the sponge surface whose maximum thickness 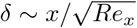, where *x* is distance along the sponge and *Re*_*x*_ = *Ux/ν*. Using the representative *x* ∼ 10 cm we obtain *δ*∼ 1 mm. Thus, the ostia through which water flows into the microfluidic network of the sponge sit within the viscous boundary layer. While the general problem of flow into a permeable wall has been studied in some detail [27–29], no connection has yet been made with sponge fluid dynamics.

The relationship between external and internal flows can then be recast in terms of the connection between Bernoulli pressure differences between the ostia and osculum set up by external flows and internal pressures created by choanocytes beating within the chambers. In this regard there are conflicting conclusions in the literature. Dahihande and Thakur [30] reported that the number density of choanocyte chambers and the number of choanocytes per chamber are positively correlated with the pumping rate of sponges. On the other hand, Larsen and Riisgård [31] found that the pumping rate depends on the pressure losses of the aquiferous system with increasing sponge size and not on the reduced density of choanocytes. Recently, there have been several important numerical studies of the sponge pump at the level of choanocytes. Asadzadeh et al. [32] found numerically that the both the glycocalyx mesh covering the upper part of the collar and secondary reticulum are important for the pump to deliver high pressure. They also reported that choanocytes arranged in a cylindrical configuration can pump water efficiently owing to the formation of a hydrodynamic gasket above the collars [33]. However, the spherical shape of choanocyte chambers was not discussed hydrodynamically in any of these former studies and the physiological significance of that shape is unclear.

Here we examine the biological significance of the body design of spherical choanocyte chambers by combining direct imaging of choanocyte chambers in living sponges with computational studies of many-flagellum models of their fluid mechanics. We used freshwater sponges *Ephydatia muelleri* as a model organism for experimental observation and to define a computational model of the fluid mechanics of a choanocyte chamber. The experimental and computational methodologies are explained in Sections II and III. In Section IV, we show computationally that the flagella beating against the flow play a role in raising the pressure inside the choanocyte chamber and the mechanical pumping efficiency reaches a maximum at a certain outlet opening angle. Section V shows the good agreement between the experimental and the numerical results. Section VI investigates the extent to which the flows in a choanocyte chamber can be represented by a small number of Stokes singularities. In the summary Section VII, we suggest further avenues of research on sponge physiology and development.

## II. EXPERIMENTAL METHODS

### Harvesting and culturing of sponges

Freshwater sponges of the species *Ephydatia muelleri*, shown in Fig. 1(a), living on natural stones were collected from the Hirose River (Izumi, Sendai City, Miyagi, Japan). The asexually produced buds of reproductive cells known as gemmules were peeled off, and the spicules on them were removed. To remove impurities, the gemmules were washed with a 1% aqueous solution of H_2_O_2_, followed by three successive rinses with pure water to remove excess H_2_O_2_. The purified gemmules were stored in a refrigerator at 4°C. Sponges were cultured in a plastic dish with Strekal’s medium at room temperature (25°C).

### Imaging

Choanocyte chambers of *E. muelleri* were imaged on an inverted microscope (IX71, Olympus Corp., Tokyo, Japan), as shown in Fig. 2(a). Flagellar beating in the chambers was imaged with a oil immersion objective (100*×*, N.A.= 1.40) and a high speed camera (500 fps, 1024*×*1024 pixels, FASTCAM SA3, Photron Limited, Tokyo, Japan) for durations of 5 s. The beating appears as a spatiotemporal brightness fluctuation in the pixels of the images [34, 35], as shown in Fig. 2(b), from which the beat frequency was measured from a fast Fourier transform (n=5, N=3). The various geometric and dynamical characteristics of choanocyte chambers (n=6-8, N=4) and the flagellar motion within them (n=3-5, N=4) are shown in Table I, in which the results of *Spongilla lacustris* obtained by Mah et al. [18] are added for comparison.

**TABLE I.** Characteristics of choanocyte chambers.

| Quantity | *E. muelleri* | *S. lacustris*<sup>a</sup> | Simulation |
| --- | --- | --- | --- |
| number of flagella $N$ | 112±31 | | 37-359 |
| flagellar length $L$ | 12.9±1.1 μm | 10.4±0.3 μm | 13 μm |
| flagellar amplitude $A$ | 1.58±0.21 μm | 3.3±0.9 μm | 0.14$L$ |
| beat frequency $\nu$ | 26.1±8.9 Hz | 11.0±1.1 Hz | |
| chamber radius $R$ | 17.4±2.5 μm | | (1.5-3)$L$ |

<sup>a</sup> Data for *Spongilla lacustris* are from Mah et al. [18].

**FIG. 2.**
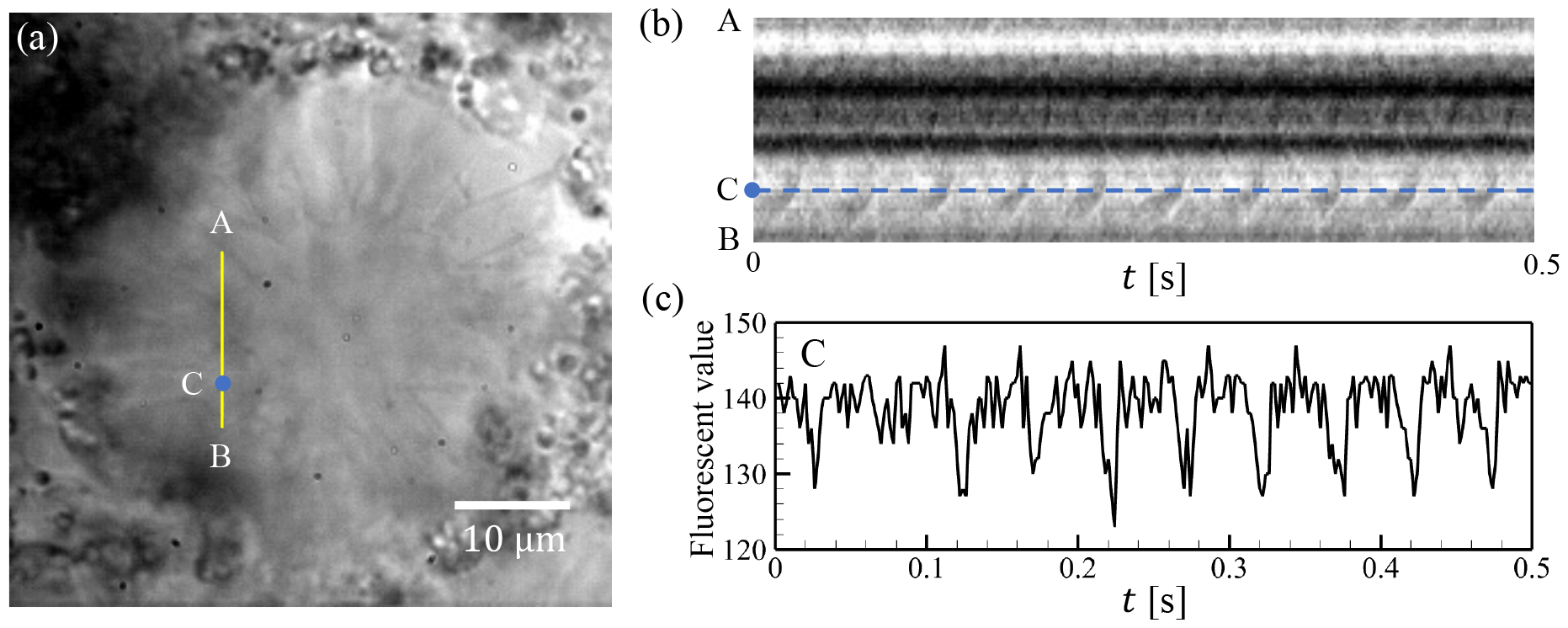
Flagellar beating in a choanocyte chamber of *E. muelleri*. (a) Microscope image of a choanocyte chamber. (b) Spaciotemporal brightness fluctuation in the choanocyte chamber showing flagellar beating. (c) Slice of data in (b) showing the temporal oscillations in pixel intensity.

## III. COMPUTATIONAL FLUID DYNAMICS OF CHOANOCYTE CHAMBERS

Choanocyte chambers consist of choanocytes arranged in a spherical configuration, each with a single flagellum oriented toward the center of the chamber, propagating bending waves radially inward. In the computations, we represent the chamber as a rigid sphere of radius *R* from which flagella emerge, as in Fig. 3. The chamber has many small inlet holes termed *prosopyles* and one large outlet hole known as the *apopyle*. Additionally, there is the gasket-like *reticulum*, a fine structure that connects the apical part of the collars, which was also observed in *Spongilla lacustris* [36]. The reticulum delivers high pressure because the choanocyte acts as a collar-vane-flagellum pump system [32]. The reticulum was modeled as a concentric spherical shell inside the choanocyte chamber (Fig. 3(b)), and its radius was varied with the chamber radius so that the length of the collar extending from the choanocyte was a constant 8.2 *μ*m [18]. Finally, *cone cells* are found in several freshwater sponges including *E. muelleri* [37], and are flattened near apopyles [36– 38]. They form a ring and connect collars with the walls of excurrent canals to prevent the disadvantageous current. The cone cell ring was modeled as a conical surface near the apopyle (Fig. 3(b)), with an opening angle *θ*_*a*_.

**FIG. 3.**
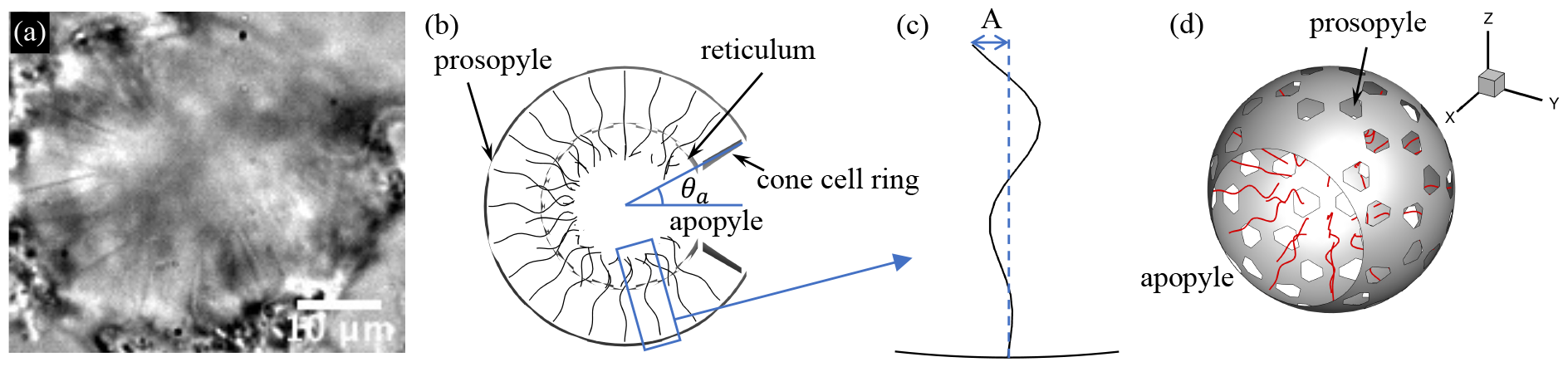
Computational model of choanocyte chamber. (a) A choanocyte chamber of *E. muelleri*. (b) Schematic cross section of the model chamber. (c) Definition of the flagellar beat amplitude *A*. (d) Three-dimensional model of the choanocyte chamber.

### Governing equations

Scaled by the flagellar motion, the Reynolds number *Re*≪ 1 in a chamber, so the fluid flow in and around choanocyte chambers is governed by the Stokes equations, and we use slender-body theory [39] for the flagella. Their centerlines are parameterized by arclength *s* ∈ [0, *L*], and we measure the chamber radius *R* in units of the flagellar length, with

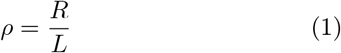

a parameter that controls the size of the central region of the chamber devoid of flagella. The velocity ***v*** at point ***x*** ∈ *s*_*i*_ located on flagellum *i* can be written as [40]

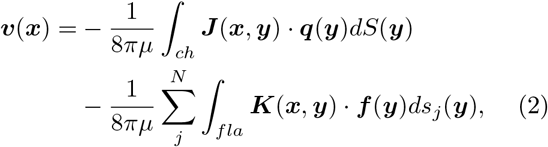

where *μ* is the viscosity, ***q*** is the viscous traction on the chamber, ***f*** is the force density of the flagella and *N* is the total number of flagella. The first integral is over all surfaces *S*, including those of the chamber, reticulum and cone cell ring. The second integral is over flagellar centerlines. In (2), ***J*** is the Green’s function

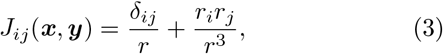

where *r* = |***r***| and ***r*** = ***x*** − ***y***, and ***K*** is the kernel [39]

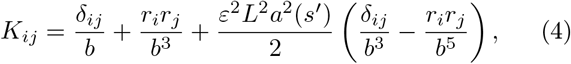

where 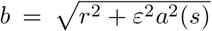. Here the radius function satisfies 0 *< a*(*s*) ≤ 1 for each 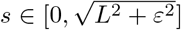, where *ε* is the ratio of the length to the radius of flagellum and is set to *ε* = 0.01. The radius function *a*(*s*) is

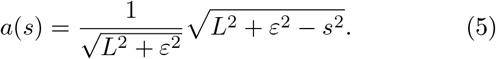

When the point ***x*** is not on a flagellum, the velocity is

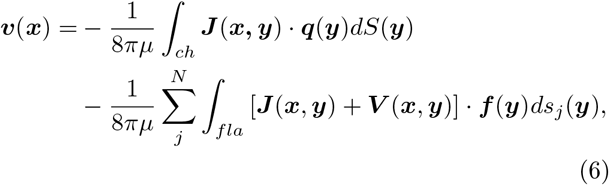

where [39, 41]

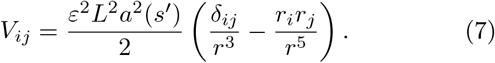

### Flagellar motions

Each of the flagella within a computational choanocyte chamber is actuated with a prescribed waveform at a frequency *ν* = 1*/T*, where *T* is the period of motion, and fixed at its base on the choanocyte chamber surface at a point ***x***_*b*_. We define the orthonormal body frame ***g***_1_ and ***g***_2 at_ ***x***_*b as*_ ***g***_1_(***x***_*b*_) = ***b***(***x***_*b*_), and ***g***_2_(***x***_*b*_) = ***b***(***x***_*b*_) ***n***(***x***_*b*_)*/* ***b***(***x***_*b*_) ***n***(***x***_*b*_), where ***b***(***x***_*b*_) = ***e***_1_ ∧ ***n***(***x***_*b*_), ***e***_1_ = (1, 0, 0) and ***n***(***x***_*b*_) is the out-ward unit normal vector to the surface. With *τ* = *t/T*, the motion of a flagellum is parameterized as

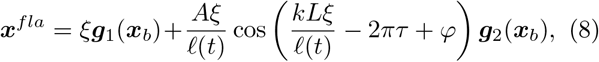

where *A* is the beat amplitude, *k* = 2*π/λ*, with *λ* the wavelength, the coordinate *ξ* ∈ [0, *ℓ*(*t*)] spans the time-dependent *projected* length *ℓ*(*t*) of the flagellum under the constraint of fixed total arclength *L*. For the oscillating flagellum described by (8), the projected arclength *ℓ* is considerably less than the total arclength *L*. For the amplitude *A* = 0.14*L* used in numerics and for the values of *kℓ* ∼ (3 − 4)*π* typical of experiment we have *ℓ/L* ∼ 0.76. This contraction plays an important role in the pressure distribution within the choanocyte chamber. As we observed no phase synchrony in our studies of sponge flagella, much as earlier studies of multicellular choanoflagellates saw no synchrony [42, 43], in computations we randomly set the phase *φ* for each flagellum in the range *φ* ∈ [0, 2*π*], reproduce the independent flagellar motion in the chamber.

### Boundary element method

When the choanocyte chamber is fixed and there is no background flow, the boundary conditions are

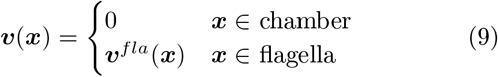

where ***v***^*fla*^ = *∂****x***^*fla*^*/∂t* is the flagella velocity, and the chamber includes the reticulum and cone cell ring.

The choanocyte chamber surface, reticulum and cone cell ring were composed of 9,710 triangular mesh elements and 5,401 nodal points. These quantities depended on the chamber radius, the size of apopyles, and the number of flagella. Each flagellum was discretized into 20 elements with 21 nodes. All physical quantities were computed at each discretized point. The boundary integral equations (2)) and (6) were computed using Gaussian integration, leading to the linear algebraic equations

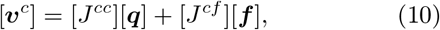

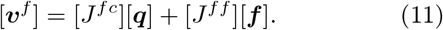

Both [***v***^*c*^] and [***q***] have size 3*N*_*c*_, and that of [***v***^*f*^] and [***f***] is 3*N*_*f*_, where *N*_*c*_ is the total number of nodes on the chamber surface, the reticulum and the cone cell ring, and *N*_*f*_ is the total number of nodes on the flagella. The matrix size of [*J*^*cc*^] is 3*N*_*c*_ *×* 3*N*_*c*_ and [*J*^*cf*^] is 3*N*_*c*_ *×* 3*N*_*f*_. Considering the boundary condition (9), the equations can be rewritten as

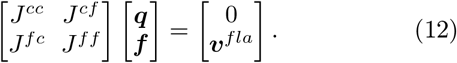

The dense matrix was solved using the lower-upper (LU) factorization technique.

### Parameters

To investigate the effects of the choanocyte chamber geometry on the pumping function of the chamber, in the numerical studies reported below we studied a range of several of the parameters of the computational modes: number of flagella *N* ∈ [37 −359], chamber radius *R* ∈ [1.5*L*, 3*L*] (with *L* = 13*μ*m), scaled wavenumber *kℓ* ∈[1.5*π*, 3*π*], and apopyle opening angle *θ*_*a*_ ∈ [10°, 90°]. Through all computations, the calculational time step *Δt* was set to 0.02*T*, where *T* is the flagellar beat period; we also used this in the time averaging method discussed below.

### Choanocyte packing fraction

In describing the numerical computations, the various parameters of the setup can be recast in convenient dimensionless quantities. First, if *θ*_*a*_ is the opening angle of the apopyle, the area of the chamber available for choanocytes is 2*πR*^2^(1 + cos *θ*_*a*_). We may view the flagellar undulation amplitude *A* in (8) as defining an area *πA*^2^ associated with each choanocyte, and thus it is natural to define the area fraction *ϕ* of choanocytes on the chamber surface as

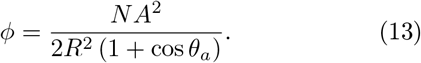

For reference, in a hexagonal close packing on a flat surface, the maximum packing fraction is *π* 3*/*6 ≃ 0.907.

## IV. NUMERICAL RESULTS

### A. Basic observations

*Flow field:* The time-averaged flow field and pressure in the center cross-section of a chamber are shown in Figs. 4(a) and (b) for representative parameters. We confirmed that the unidirectional flow, from prosopyle to apopyle, is generated by flagellar beating in the spherical chamber. In this spherical geometry, the pressure field can be divided into low and high pressure regions, as previously reported [32, 36]. The low pressure region sucks water from the prosopyle, while the high pressure region ejects water outward, leading to a unidirectional flow with volumetric rate *Q*^*^ from the apopyle of

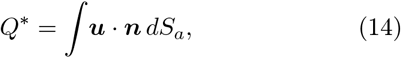

where *S*_*a*_ is the area of the curved surface of the apopyle, ***u*** is the velocity on *S*_*a*_, and ***n*** is the outward unit normal vector on *S*_*a*_. As the flagellar length is the fixed scale used for nondimensionalizing quantities, we use it and the beat period to define the rescaled flow rate

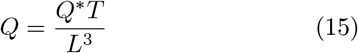

and its time-averaged value 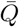. As shown in Fig. 4(c), *Q* does not vary significantly with time. For the parameters *ρ* = 1.5, *N* = 143 (*ϕ* = 0.33), *θ*_*a*_ = 30° and *kℓ* = 3*π* we find 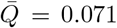, a value that is compared with experimental results in Section V.

**FIG. 4.**
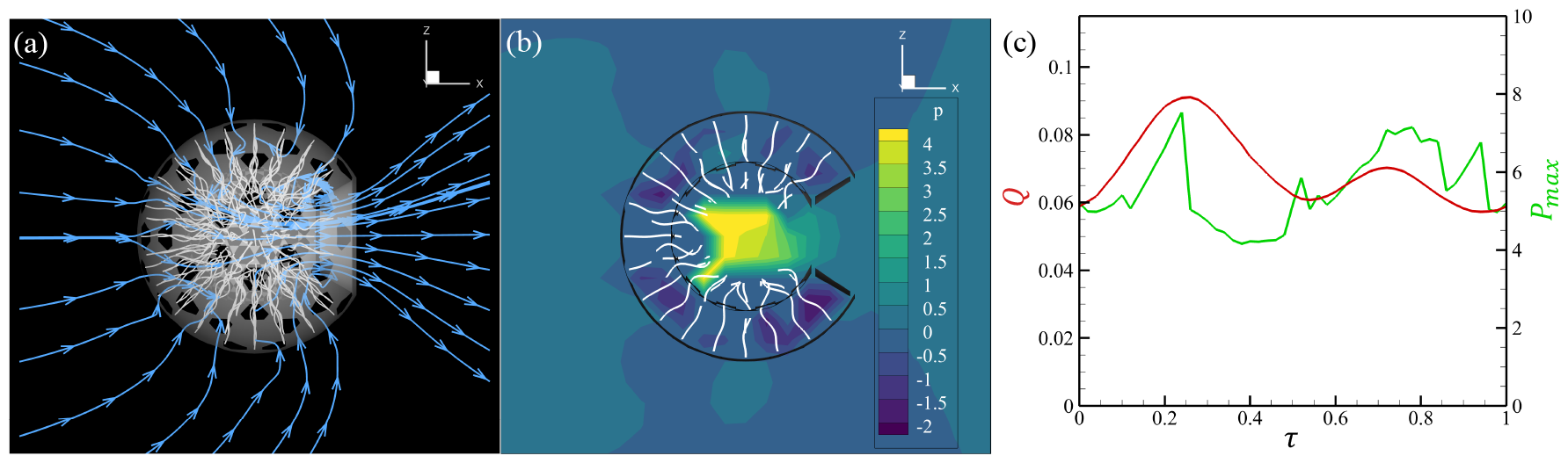
Numerical results. (a) Flow field around the model choanocyte chamber, with *ρ* = 1.5, *N* = 143 (*ϕ* = 0.33), *θ*_*a*_ = 30° and *kℓ* = 3*π*. The streamlines are drawn in the center cross-section. (b) Pressure distribution in the chamber. (c) Temporal behavior of the outlet flow rate and maximum pressure within the chamber.

The pressure at point ***x*** is [44, 45]

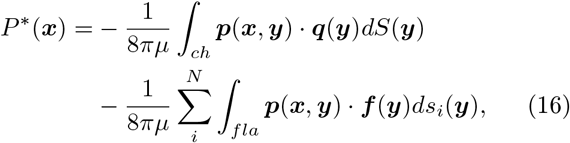

where ***p*** = 2***r****/r*^3^. From the Stokesian balance **∇***P*^*^ = *μ*∇ **u**, with *u* ∼ *L/T*, a suitable rescaled pressure is

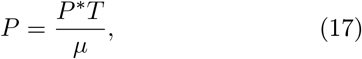

which is used to quantify the maximum pressure *P*_max_ and its time-average 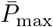. The time course of *P*_max_ is shown in Fig. 4(c), in which the apparent discontinuities arise from abrupt changes in the location of the maxima due to the random flagellar phases.

*Flagellar force:* The force density ***f*** and the velocity ***u*** of node points on a flagellum are shown in Fig. 5(a). These can be used to determine the rate of working *W*^*^ of the entire flagellum in moving fluid as

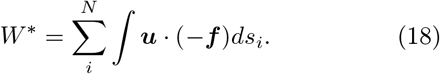

**FIG. 5.**
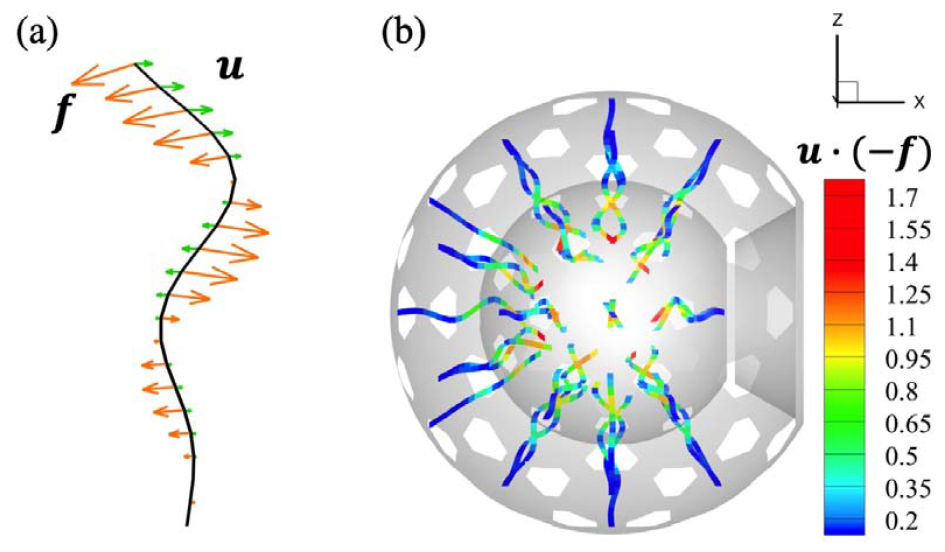
Work rate of flagella. (a) Force density and velocity at node points of a flagellum. (b) Magnitude of the inner product of the force density and the velocity.

With the force density ***f***∼ *μ****u*** and speeds scaling as *L/T*, we define the dimensionless rate as

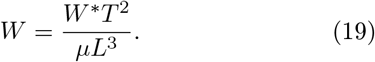

Intuitively, we see that the magnitude of ***u*** (−***f***) of each flagellum is largest near the tip, as in Fig. 5(b), which creates high pressure in the center of the chamber.

### B. Output increase with number of flagella

To clarify the relationship between the number of flagella in a chamber and its fluid dynamical properties, pumping functions were investigated as the number was varied from 10 to 347. The pumping function was evaluated according to the outlet flow rate *Q* from the apopyles, the rate of working of the flagella, the maximum pressure *P*_*max*_ in the chamber (assuming zero pressure at infinity) and the mechanical pumping efficiency *η* averaged over time during a beat cycle of the flagellar, where

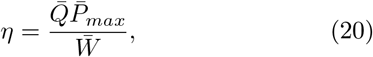

where overbars indicate time-averaged quantities.

In Fig. 6, ensemble averaged 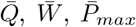, and *η* are shown with three independent phases and rotation angles of the flagellar beating planes. The larger the number of flagella (and the greater the packing fraction *ϕ*), the greater 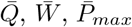, and *η*, indicating better performance as a pump. Most notably, the average flux 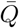 is very nearly linear in *N* and the flux per flagellum 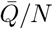 saturates to a constant value of 5 × 10^−4^ at large *N*. In interpreting this value it is useful to adopt a simplified view of the flagellum as a localized point force acting on the fluid, a model that has experimental support from studies of *Chlamydomonas reinhardtii* [46]. The associated Stokeslet flow field, if examined in free space without boundaries, has an infinite flux through any plane orthogonal to the direction of the point force, while for a force orthogonal to a nearby no-slip surface the flux vanishes by fluid conservation. This latter result arises from compensating flows away from the surface near the singularity and toward the surface far away from it. A numerical computation of the flux associated only with the outgoing flows driven by a single model flagellum used in the present calculations, attached to a no-slip wall, gives a value of∼ 0.01, a factor of 20 greater than the limit seen in Fig. 6(a). While the porosity of the wall would tend to increase the flux, the tight confinement within the chamber and the typically small apopyle angle clearly leads to significant cancellation. For a given chamber radius, excluded volume effects from cells and their collars limits the number of choanocytes that can be spherically aligned, a limit that depends on the size and shape of the choanocyte. The choanocyte diameter varies among species: 3 *μ*m for *Tetilla serica* [47], 3.7 5 *μ*m with pseudocylindrical shape for *Halisarca dujardini* [48], 6 *μ*m for *E. fluviatilis* [49] and 5 *μ*m for *E. muelleri* [50].

**FIG. 6.**
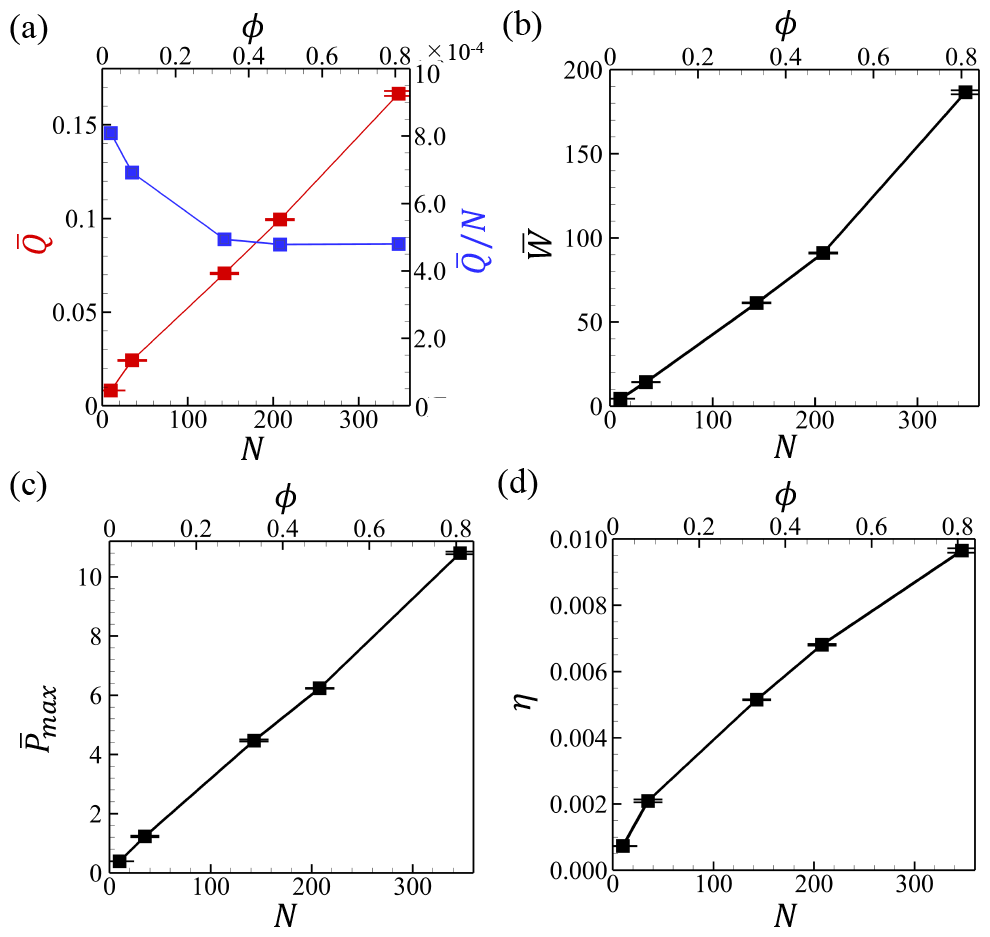
Correlation of pumping functions with the number of flagella. (a) Outlet flow rate. (b) Work rate done by flag-ella. (c) Maximum pressure in the chamber. (d) Mechanical pumping efficiency.

### C. A smaller chamber radius increases efficiency

We investigated the effect of the chamber radius on the pumping functions such as the efficiency by varying the radius from 1.5*L* to 3.0*L* while keeping the number density of flagella fixed. As shown in Fig. 7, the flow rate and work rate increased with the chamber radius as the number of flagella also increased. On the other hand, the maximum pressure decreased as the chamber radius increased. The pressure drop was due to the large center space within the chamber, where there are no direct flagella forces. As a result, the mechanical pumping efficiency decreased with increasing radius. These results indicate that a larger chamber has no efficiency advantage, but rather that it is advantageous to pack more choanocytes into a smaller chamber.

**FIG. 7.**
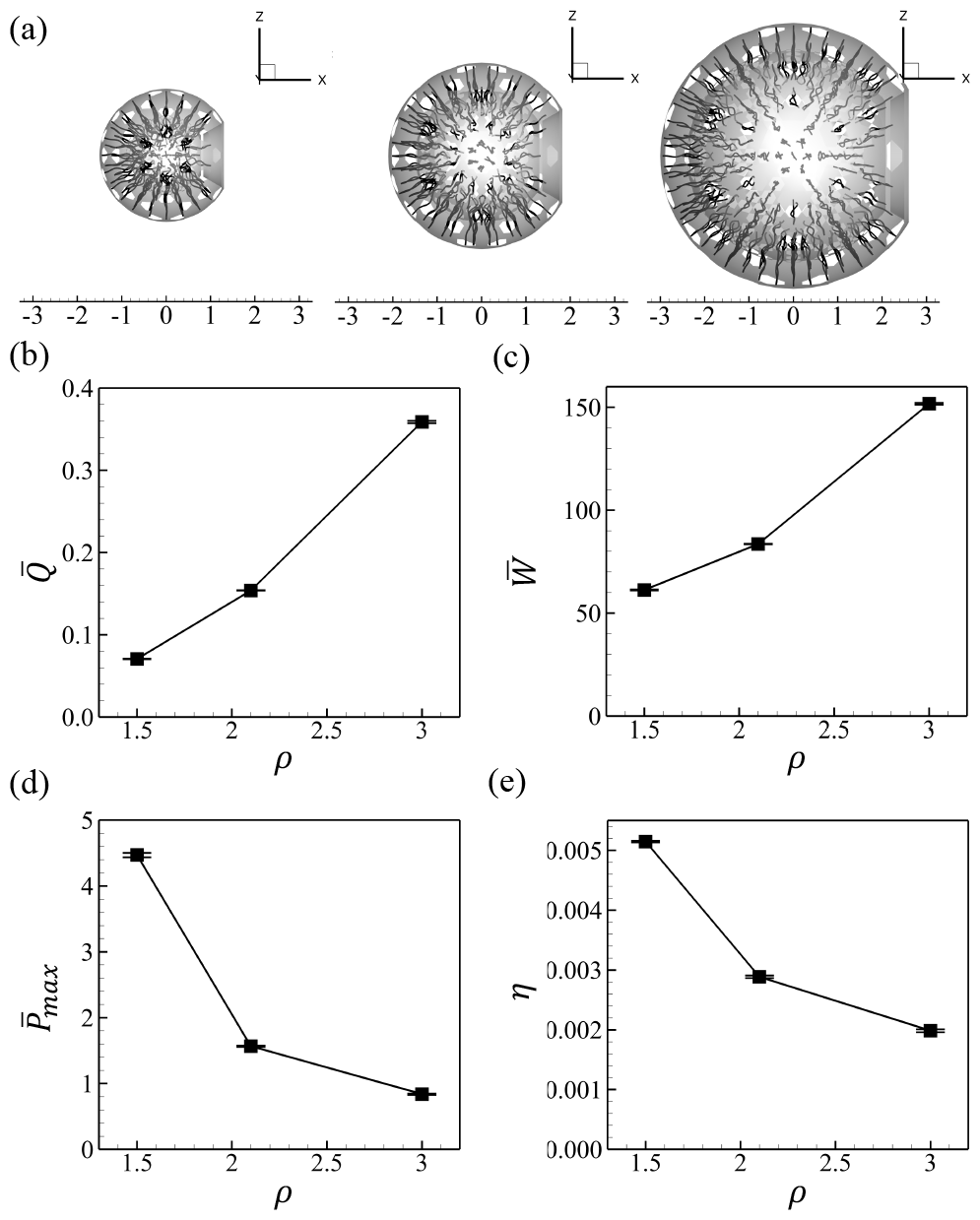
Correlation of pumping function with chamber radius. The length and number density of flagella are fixed. (a) Outlet flow rate. (b) Work rate done by flagella. (c) Maximum pressure in the chamber. (d) Mechanical pumping efficiency.

### D. Efficiency is maximized at intermediate *kℓ*

We studied the effect of changing the flagellar wave number while keeping the flagellar length fixed. Interestingly, as shown in Fig. 8, the outlet flow rate and mechanical pumping efficiency exhibit peaks at intermediate values of *k*, while the particular value of the peak *k* differs between the two. The mechanical pumping efficiency reaches a maximum when at the relatively low wave number *kℓ* = 3*π*, where *ℓ* is projected flagellar length. That higher wave numbers lead to reduced efficiency, despite the reduced work rate of the flagella, arises from an effect similar to that found when the chamber radius is reduced; Since *L* is fixed, *ℓ* shrinks at higher wave numbers, increasing the space at the chamber center from which flagella are absent, and the central pressure reduces with higher wave number. The wave number associated with the efficiency is maximized is compared with the experimental results in Section V.

**FIG. 8.**
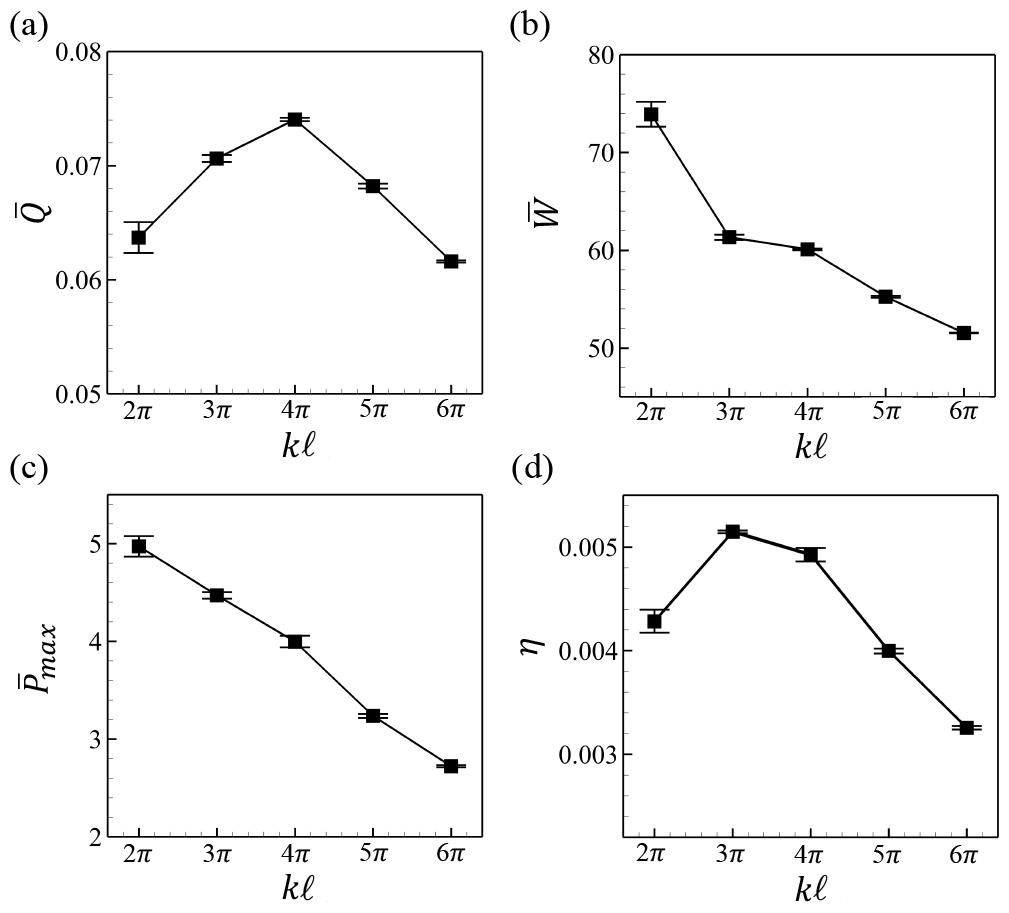
Correlation between the pumping function and wave number of flagella. (a) Outlet flow rate. (b) Work rate done by flagella. (c) Maximum pressure in the chamber. (d) Mechanical pumping efficiency.

### E. Efficiency is maximized at intermediate *θ*_*a*_

While the choanocyte chamber diameter, apopyle area, and diameter were examined in previous studies [22, 51, 52], it has remained unclear how the apopyle’s aperture ratio affects the pumping function of the chamber. We examined the pumping function *θ*_*a*_ ∈ [10°, 90°] while keeping the area fraction *ϕ* within each of three narrow ranges, adjusting the number of flagella with *θ*_*a*_ accordingly. Figure 9(b-e) shows the variation in pumping functions with *θ*_*a*_. With all three flagella densities, similar trends were observed, with small variations in the position of various peaks. The outlet flow rates reaches a maximum at *θ*_*a*_∼ 40° − 60°. On the other hand, the maximum pressure decreases monotonically with *θ*_*a*_, an effect that arises from the reduction in the the number of flagella directed against the bulk flow as *θ*_*a*_ increases. Thus, the spherical shape of the choanocyte chamber has the effect of increasing pressure. Since the pumping efficiency is the product of the maximum pressure and the flow rate, its peak in Fig. 9(e) shifts to the lower *θ*_*a*_ regime of 20°− 50° compared to the case of the flow rate (cf. Fig. 9(b)). From these results, we conclude that flagella around apopyles, which seem to disturb unidirectional flow, contribute to creating the high pressure rise and that the choanocytes with intermediate but small *θ*_*a*_ can achieve high mechanical pumping efficiency.

**FIG. 9.**
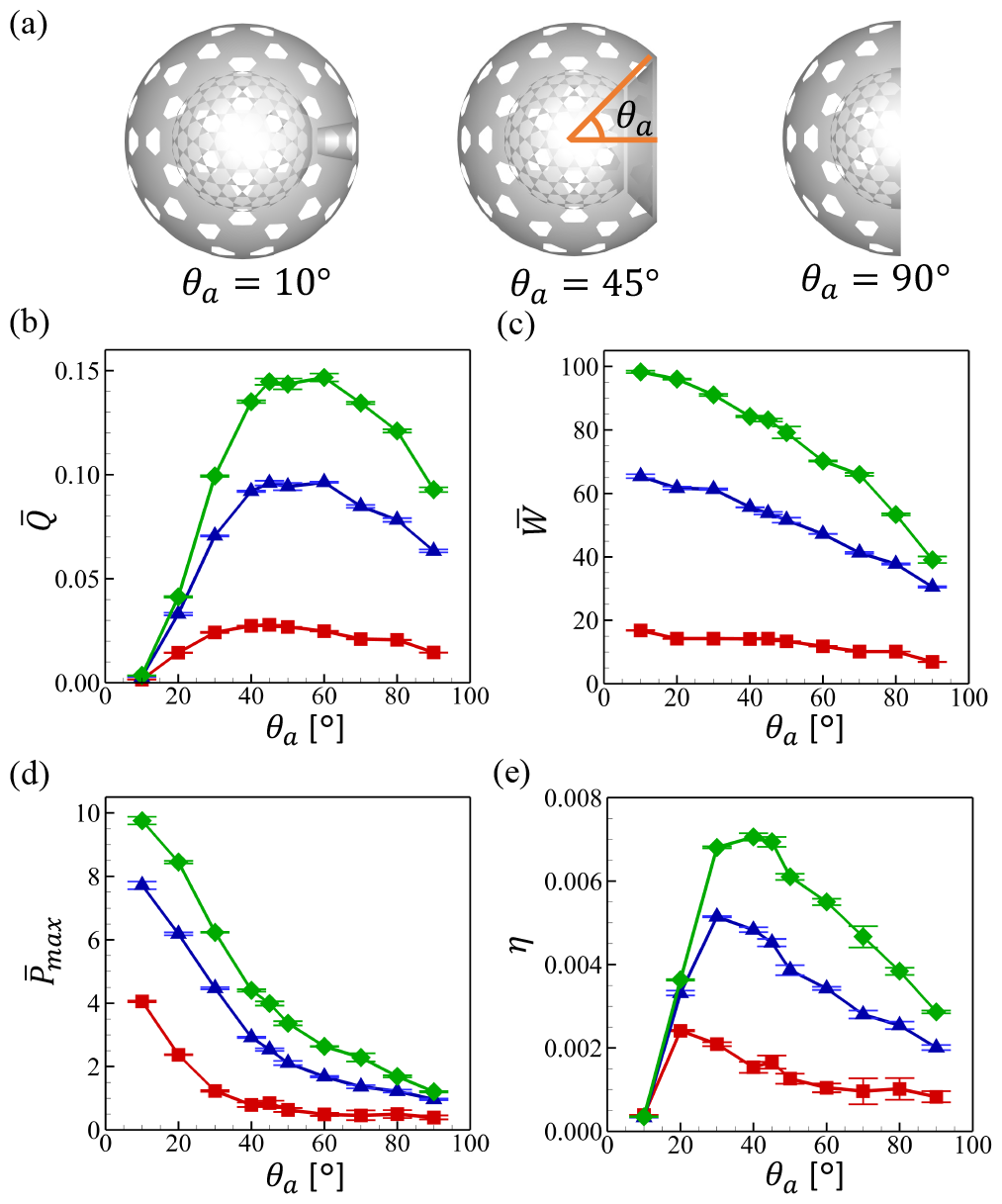
Correlation between pumping function and outlet opening angle. (a) Computational domain for three values of *θ*_*a*_. (b-e) Chamber properties as a function of opening angle, keeping area fraction *ϕ* in narrow ranges: red: 0.074 − 0.093, blue: 0.31 − 0.34, and green: 0.39 − 0.49. (b) Outlet flow rate. (c) Work rate of flagella. (d) Maximum pressure in the chamber. (e) Mechanical pumping efficiency.

## V. COMPARISON WITH EXPERIMENT

### Pumping flow rate

The numerical results in Section IV A, with parameters taken from experiment, yielded time-averaged flow rates of a choanocyte chamber of 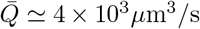. This can be compared with previous experimental results. The flow rate of the choanocyte chamber of *Haliclona urceolus* was estimated as *Q* = 4.5 *×*10^3^*μ*m^3^*/*s [22], which is very close to the present results. In addition, the flow rate per choanocyte of a syconoid type sponge was estimated at 48 *μ*m^3^/s [33], and if the flow rate scales with the number of choanocytes, estimated to be 100 (cf. Table I), we obtain *Q* = 4.8 *×* 10^3^*μ*m^3^/s, also in good agreement with our results. Thus, the flow rate obtained in this study is consistent with former experimental observations.

### Flagellar wave number

Next, we compare the wave number at which the efficiency is maximized with our experimental results. The mechanical pumping efficiency as a function of the wave number with three different number densities of flagella is shown in Fig. 10(a). We see peaks in efficiency with the *kℓ*∼ (3 − 4)*π*, independent of the number density of flagella. In our experiment, the flagellar motion of *E. muelleri* choanocyte was extracted from the high-speed imaging as shown in Fig. 10(b). From the projected flagellar length *ℓ* = 9.83 *±* 0.74 *μ*m and the wavelength *λ* = 5.98 *±* 0.48 *μ*m, we find *kℓ* = (3.30 *±* 0.27)*π*, in good agreement with the numerical results for the maximum efficiency. Similar trends are seen in other species. In the case of *S. lacustris*, for example, *kℓ* ∼ 4*π* using *ℓ* = 10.4 *μ*m and *λ* = 5 *μ*m [18, 32]. Thus, the flagellar wave number appears to be tuned to maximize the mechanical pumping efficiency.

**FIG. 10.**
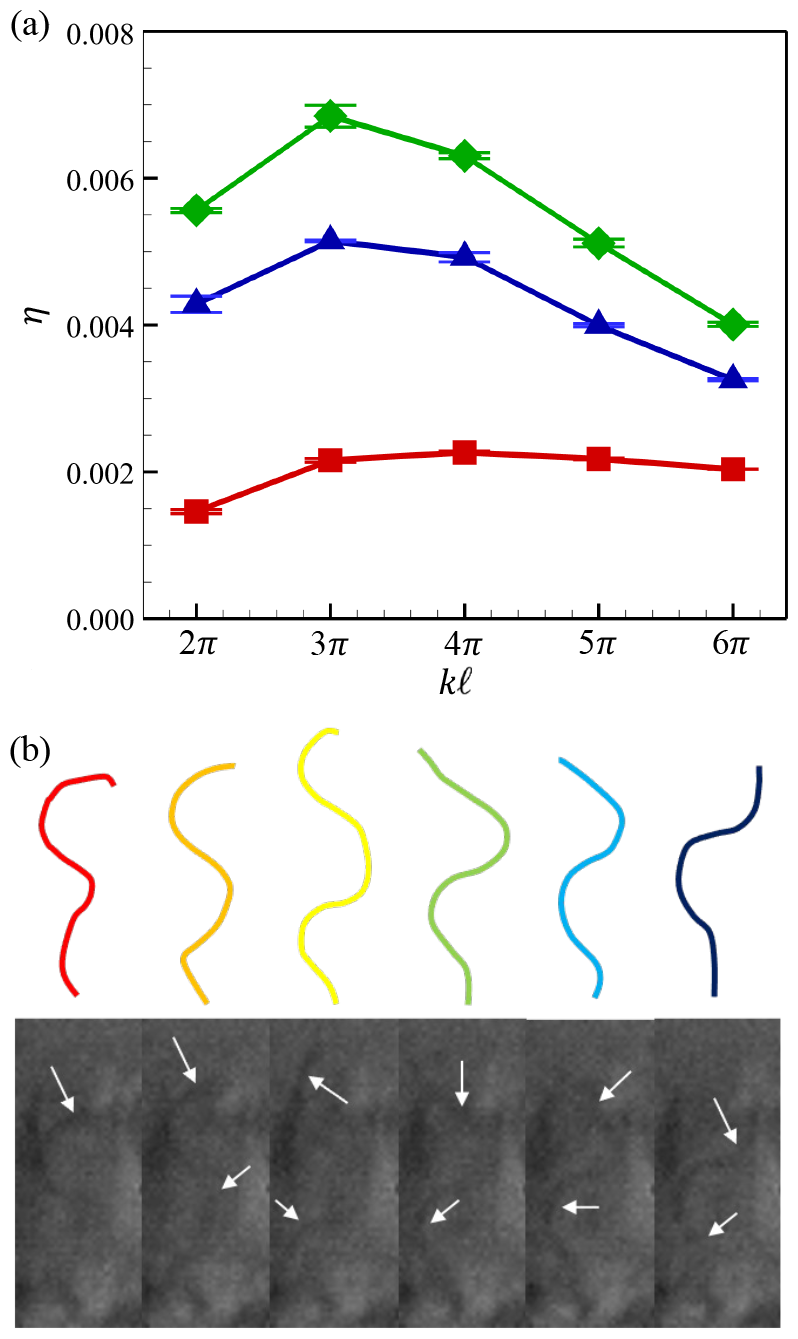
Wave number of flagella. (a) Mechanical pumping efficiency as a function of the wave number with color scheme as in Fig. 9 for *ϕ* in the ranges 0.074 − 0.093 (red), 0.31 − 0.34 (blue), and 0.39 − 0.49 (green). (b) Flagellar motion of a choanocyte of *E. muelleri*. White arrows indicate flagella. Colored curves represent extracted flagellar waveforms.

### Chamber size and choanocyte density

Our experimental observations show that the number of flagella in a choanocyte chamber of *E. muelleri* is ∼ 100, similar to the value ∼ 80 found in earlier work on choanocytes of *H. urceolus*, in which the chamber diameter is ∼ 30 *μ*m [22]. In several marine sponges, the chamber diameters ranges from 16 − 31 *μ*m and the number of choanocytes ranges from 32− 130, which indicates that choanocytes are densely packed in the chamber. There is a positive correlation between the number of choanocytes per chamber and the pumping rate of a sponge according to experimental observations [30]. Hence, it is inferred that the chamber is filled almost completely with as many choanocytes as allowed by the chamber size. These tendencies are consistent with our finding that larger number of flagella enhances pumping function and efficiency.

As noted earlier, there is an upper limit on the number density of choanocytes due to the excluded volume of cells and the aperture ratio of apopyles and prosopyles. Hence, to increase the number of choanocytes, the choanocyte chamber is forced to make its radius larger. On the other hand, when the radius is large, the pumping efficiency becomes low. These two conditions are contradictory, so a balance must be reached between them. To find balanced conditions in nature, we plotted the correlation of the flagellar length and the choanocyte chamber diameter for various species of sponge as shown in Fig. 11. We see an obvious positive correlation between them, indicating that the chamber diameter increases with flagellar length. We analytically calculated the hypothetical minimum diameter of *E. muelleri* under the condition that adjacent flagella of amplitude 3.16 *μ*m do not contact each other. The result is 36.4 *μ*m, which is close to the actual diameter of 34.7 *μ*m. This implies that the chamber diameter is designed to be as small as possible while avoiding overlapping flagella. Smaller chamber diameters may have other advantages, such as the ability to accommodate a larger number of cells in a smaller volume and greater flexibility in the design of the canal in the entire of sponge body.

**FIG. 11.**
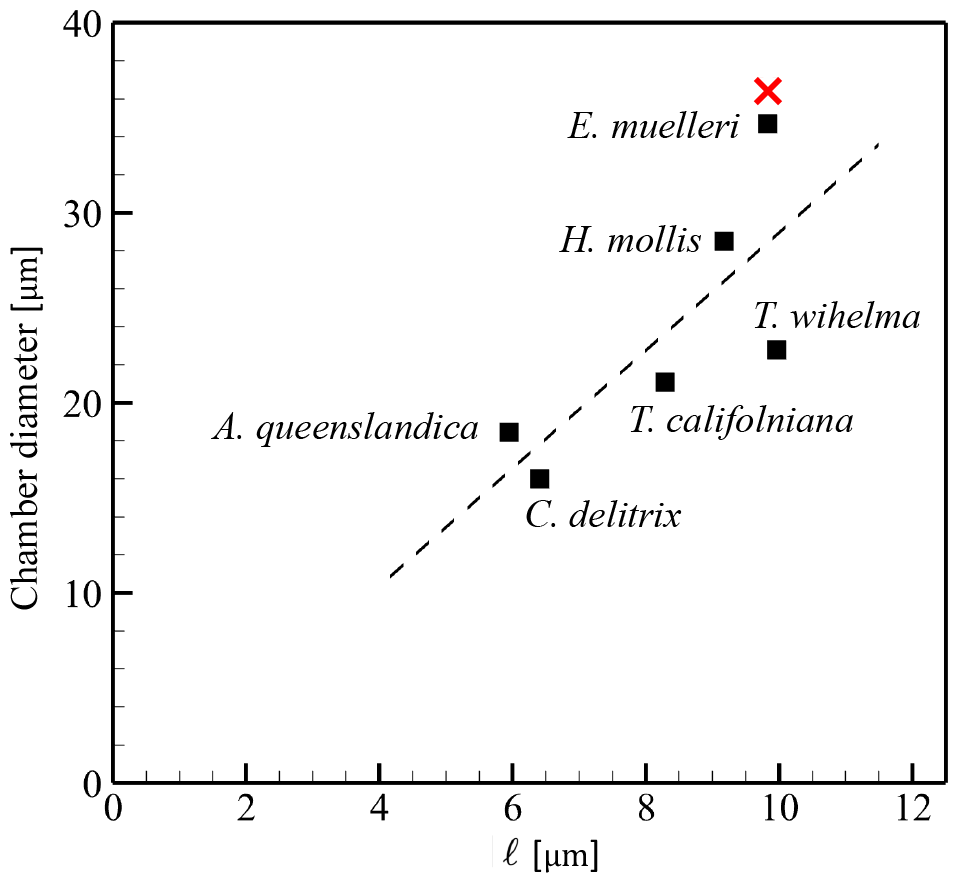
Correlation between the projected flagellar length and the choanocyte chamber diameter. If not specified, the values were measured manually from images; *Tethya wilhelma* from Hammel et al. [25]; *Haliclona mollis, Tathya californiana*, and *Cliona delitrix* from Ludeman et al. [24]; *Amphimedon queenslandica* from Sogabe et al. [53]. *×* indicates the hypothetical minimum diameter of *E. muelleri* that is obtained under the condition that adjacent flagella do not touch.

### Outlet opening angle

Finally, we compare the experimentally observed opening angles with the range *θ*_*a*_ = 20°−50° found to be maximal for pumping efficiency. We observed several cross-sections of a choanocyte chamber of *E. muelleri* by shifting the focal plane as shown in Fig. 1(c-f). From the image around the apopyle (cf. the top focal plane in Fig. 1(e)), we measure the apopyle diameter. Using this value and the chamber diameter, and assuming a spherical chamber shape, the average apopyle opening angle was found to be be *θ*_*a*_ = 32.2°. In the case of *E. muelleri*, the number density of flagella is 0.032*/μ*m^2^, which, with the chamber diameter of 34.7*μ*m, and the flagella number of 112 gives a coverage fraction *ϕ* = 0.25, associated with the blue data in Fig. 9(e) for pumping efficiency, which is maximized at *θ* = 30°. The very good agreement with the observed value for *E. muelleri* indicates that the natural configuration of the choanocyte chamber of *E. muelleri* optimizes the mechanical pumping efficiency.

The geometric properties of choanocyte chambers in many sponge species that have been reported previously are collected in Table II. To compare the present study with these former ones, we plotted the opening angle *θ*_*a*_ versus the number density of choanocytes in a chamber. In estimating these values, we assumed the chamber is spherical. Studies on *E. muelleri, Haliclona urceolus, Haliclona permollis, Aphocallistes vastus, Neopetrosia problematica, Haliclona mollis, Tethya californiana, Callyspongia vaginalis*, and *Cliona delitrix* [22, 24] explicitly reported information about apopyles, so we could fully incorporate the effect of apopyles when calculating the number density of choanocytes. In studies on *Amorphinopsis foetida, Callyspongia* sp., *Haliclona* sp. and *Ircinia fusca* [30], on the other hand, the apopyle area was not reported and we calculated the number density without taking apopyles into account, thus underestimating the number density. The results are plotted in Fig. 12, in which the present results on those values of *θ*_*a*_ that correspond to the maximum mechanical pumping efficiency at various densities are indicated in red. The opening angles of natural sponge choanocyte chambers are clearly close to the computational results for those with maximum efficiency, thus demonstrating that many species of sponge have choanocyte chambers that, like *E. muelleri*, can achieve high mechanical pumping efficiency.

**TABLE II.** Dimensions of choanocyte chambers.

| | Chamber diameter<br>($\mu\text{m}$) | Apopyle diameter<br>($\mu\text{m}$) | Apopyle area<br>($\mu\text{m}^2$) | $\theta_a$ ($^\circ$) | Choanocytes<br>per chamber |
| --- | --- | --- | --- | --- | --- |
| 1. *Ephydatia muelleri*<sup>a</sup> | 34.7±5.0 | 18.1±4.3 | | 32.2±6.7 | 112±31 |
| 2. *Haliclona urceolus*<sup>b</sup> | 30 | 14 |  | 27.8 | 80 |
| 3. *Haliclona permollis*<sup>c</sup> | 30 | 14 |  | 27.8 | 95 |
| 4. *Aphrocallistes vastus*<sup>c</sup> | 56 | 26 |  | 27.7 | 260 |
| 5. *Neopetrosia problematica*<sup>c</sup> | 23.3 | 16.0 |  | 43.4 | 80 |
| 6. *Haliclona mollis*<sup>c</sup> | 28.5 | 14.1 |  | 29.7 | 139 |
| 7. *Tethya californiana*<sup>c</sup> | 21.1 | 0.90 |  | 2.44 | 99 |
| 8. *Calyspongia vaginalis*<sup>c</sup> | 19.7 | 5.97 |  | 17.6 | 93 |
| 9. *Cliona delitrix*<sup>c</sup> | 16.0 | 4.23 |  | 15.3 | 50 |
| 10. *Amorphinopsis foetida*<sup>d</sup> | 17.46±0.13 | | 124.11 | 46.1 | 84.11±3.02 |
| 11. *Calyspongia* sp.<sup>d</sup> | 19.35±0.2 | | 168.54 | 49.2 | 121.11±4.33 |
| 12. *Haliclona* sp.<sup>d</sup> | 19.69±0.23 | | 113.21 | 37.6 | 68.83±2.82 |
| 13. *Ircinia fusca*<sup>d</sup> | 30.68±0.29 | | 211.25 | 32.3 | 120.35±8.98 |

<sup>a</sup> $n = 5$ , $N = 3$ <sup>b</sup> Data from Larsen et al. [22].<sup>c</sup> Data from Ludeman et al. [24].<sup>d</sup> Data from Dahihande et al. [30].

**FIG. 12.**
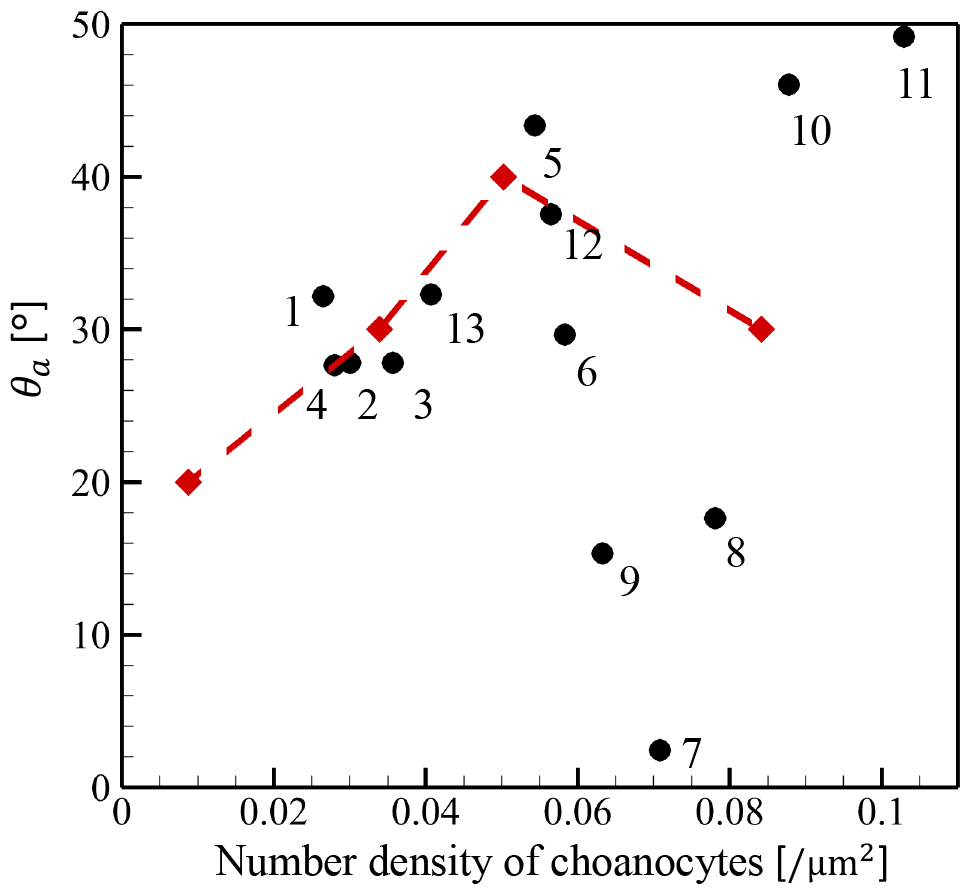
Correlation of the opening angles of apopyles and the number density of choanocyte. Black symbols and numbers correspond to the species listed in Table II. Red symbols indicate the angle at which the numerically obtained mechanical pumping efficiency reaches its maximum at each of four number densities used in the simulation.

## VI. POINT FORCE MODEL

A recurring theme in biological fluid dynamics is the representation of flow fields around multiflagellated organisms by suitable singularities in Stokes flow. For example, experimental studies of freely-swimming colonies of the green alga *Volvox carteri*, a spheroid consisting of ∼ 10^3^ biflagellated cells on its surface, have shown a dominant far-field behavior associated with a Stokeslet arising from the density offset between the colony and the surrounding water; a single point force accurately summarizes the effects of a thousand cilia [54]. At smaller scales, an accurate representation of the swirling flows near a single biflagellated alga *Chlamydomonas reinhardtii* when it swims in a breaststroke fashion requires three point forces: one for the cell body and one each for the opposing flagella [54, 55].

Returning to the densely-packing choanocyte chambers, it is natural to examine the extent to which the fluid dynamical properties we have found in the numerical studies described above can be represented by the action of one or several point forces. Such a representation would be useful in understanding the input-output characteristics of the chamber, particularly within a coarse-grained model of the the sponge network.

In the original full simulation, there were flagella, the reticulum and the cone cell ring inside a spherical chamber, and the flow was generated as shown in Fig. 13(a). We then integrated the forces acting on these internal structures consolidated them into a set of point forces. Fig. 13(b-d) shows the flow field when the internal structure is divided almost equally into *M* sections and *M* point forces are applied at the indicated locations, for the cases (b) *M* = 35, (c) 10 and (d) 1 point force at the sphere center. In all of these coarse-grained models, a flow can be observed from the center of the spherical chamber towards the apopyle.

**FIG. 13.**
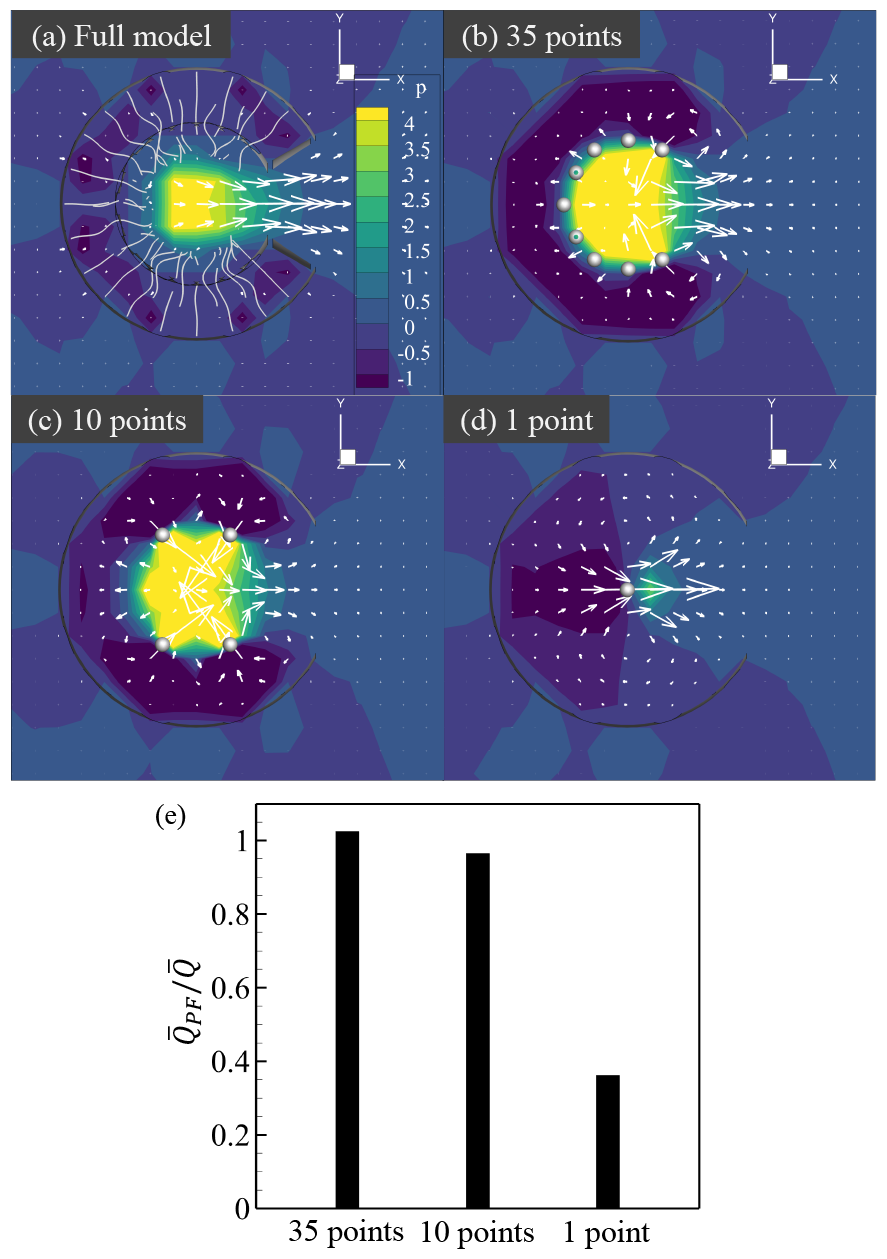
Flow generated by coarse-grained models of a choanocyte chamber. (a-d) Flow and pressure fields generated by the full simulation (a), 35 point forces (b), 10 point forces (c), and a single point force (d), where dots indicate the positions of the point forces. (e) Flow rate of the coarse-grained models 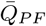 relative to the flow rate of the full simulation 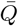.

Fig. 13(a-d) also show the pressure field. Due to the different distribution of forces, the pressure distribution is also altered from one value of *M* to another, but for all values *M >* 1 there is a similar scale of the high pressure acting at the centre of the chamber. It is only when *M* = 1 that there is a significant reduction in the central pressure. This can be seen in the flow rate of the coarse-grained models relative to that in the fully-resolved numerics, shown in Fig. 13(e). Even for *M* as small as 10 flux is quantitatively matched within a few percent. But the single point force drastically underestimates the flux as a direct consequence of underestimating the central pressure. We conclude that the effects of flagella around the apopyle, opposed to the directional flow, are indeed crucial to obtain the scale of flux from choanocyte chambers seen in experiment. At the same time, a significantly simplified representation of the chamber is possible.

## VII. CONCLUSIONS

In this study, we combined direct imaging of choanocyte chambers in living sponges with computational studies of many-flagellum models of their fluid mechanics to unravel the biological significance of the spherical shape of choanocyte chambers. We determined that there are ideal conditions for achieving maximum mechanical pumping efficiency for such a geometry. First, to achieve better efficiency, the chamber radius should be as small as possible and the number density of choanocytes per chamber should be as large as possible. Excluded volume effects limit the range of densities, and we found for *E. muelleri* and several other species that the chamber diameter is as small as possible while avoiding overlapping flagella. Second, the optimum wave number for mechanical pumping efficiency was found to be *kℓ* ∼ 3*π* when the chamber radius is 1.5 times the flagellar length. This linkage was in good agreement with experimental results on *E. muelleri*. Third, the optimum opening angle for mechanical pumping efficiency was found to be in the range 20°− 50°, with a value for *E. muelleri* of 30°, which was again in good agreement with the experimental result of 32.2°. The computational estimate of the optimum opening angle is also in good agreement with those of many other species.

In addition, our computational analysis revealed that those flagella that beat against the dominant flow play a role in raising the pressure inside the choanocyte chamber. As a result, the mechanical pumping efficiency—calculated from the pressure rise and flow rate—reaches a maximum at a modest outlet opening angle. Given that high pumping efficiency is achieved in many species (cf. Fig. 12), the choanocyte chamber may have evolved to optimize this feature, which can be viewed as a means of overcoming the high fluid dynamical resistance of the complex canal. Finally, out investigation of the fidelity of coarse-grained models of the chamber reveals that such simplifications can be taken too far to be accurate, particularly when much of the phenomena of interest are in the near-field regime, as they appear to be for sponge choanocyte chambers. This result does not preclude appropriate coarse-grained models for networks of chambers, only that the input-output characteristics are nontrivial. Logical next steps include development of such models, and on a more fundamental level understanding the developmental processes that lead to such complex network architecture in leuconoid sponges.

## ACKNOWLEDGMENTS

This work was supported by JST SPRING, Grant Number JPMJSP2114, JST PRESTO (Grant Number JPMJPR2142). K.K. was supported by the Japan Society for the Promotion of Science Grant-in-Aid for Scientific Research (21H05303, 21H05306, 22H01394) and JST FOREST Program, Grant Number JPMJFR2024. T.I. was supported by the Japan Society for the Promotion of Science Grant-in-Aid for Scientific Research (JSPS KAKENHI Grant No. 21H04999 and 21H05308). R.E.G. acknowledges support from the Wellcome Trust Investigator Grant 207510/Z/17/Z and The John Templeton Foundation.

